# Mutation and Recombination Rates in Free-Ranging Polar Bears

**DOI:** 10.64898/2026.11.05.764675

**Authors:** D Desai, LR Rivkin, L Newediuk, JM Northrup, CJ Garroway, ES Richardson, ABA Shafer

## Abstract

Mutation rate (μ) and recombination rate (r) are fundamental population-genetic parameters, yet demographic and conservation inference in polar bears (*Ursus maritimus*) has relied on values borrowed from humans and other mammals. Known values vary by an order of magnitude across mammals, and inaccuracies in these estimates propagate into downstream analyses, such as inferences of demographic history. To address this, we used high-coverage whole-genome resequencing of eight parent-offspring trios from the Western Hudson Bay subpopulation of polar bears. We identified 329 de novo mutations and estimated a pedigree-based germline mutation rate of 1.22 x 10⁻⁸ per site per generation, which is within the mammalian range and close to values previously assumed for this species. Trio-based phasing yielded 522 crossovers across 16 meioses (32.6 per meiosis), and a pedigree-inferred recombination rate of 1.50 cM/Mb. From 44 unrelated individuals sampled from the same subpopulation, we inferred a demography-aware recombination map with a length-weighted genome-wide recombination rate of 3.03 cM/Mb (median 1.28 cM/Mb). We also identified 8,515 recombination hotspot regions covering 2.10% of the genome, accounting for 43.8% of total map length, a 20.9-fold concentration. Crossovers overlapped LD-defined hotspots 5.06-fold more often than expected by chance (p < 0.001). Nucleotide diversity was low and positively rank-correlated with recombination rate, consistent with linked selection. Genomic features explained only ∼3% of recombination rate variation, with GC content and SINE abundance the only significant predictors. These species-specific parameters provide an empirical foundation for future demographic reconstruction and population projection in this vulnerable, ice-obligate species.

## 1. Introduction

Mutation and recombination are key evolutionary forces. The rates at which they occur, mutation rate (µ) and recombination rate (ρ), are also fundamental population genetics parameters. By generating novel alleles, mutations result in genomic innovation and evolutionary adaptation, but also inherited diseases; accordingly, the mutation rate dictates a species’ evolutionary pace (Rahbari et al. 2016; Bergeron et al. 2021; Gong et al. 2021). Genetic recombination (i.e. crossing-over) in meiosis produces novel recombinant gametes in sexually reproducing organisms (Hinch et al. 2019), and similarly, the rate of recombination across the genome influences the efficacy of selection (i.e., breaking linkage disequilibrium [LD]), facilitates adaptation to changing environments by generating novel genotypes, and can influence overall genetic diversity within populations (Stapley et al. 2017). Together, µ and ρ shape genetic diversity and facilitate adaptation, and as such, estimates of both parameters are usually required for demographic inferences (Hinch et al. 2019; Bergeron et al. 2021; Suárez-Menéndez et al. 2023).

In population genetic models, µ impacts the branch length in phylogenies, scales demographic estimates in coalescent models (Paradis 2016; Schiffels and Wang 2020), and is needed to convert model-derived estimates into actual quantities, such as years and abundance (Suárez-Menéndez et al. 2023). Specifically, in coalescent models, times and rates are scaled using *μ* and generation times to derive effective population sizes (*N_e_*) (Schiffels and Wang 2020). Population-scaled recombination rate (*ρ*) is related to the per-generation recombination rate by *ρ* = 2 · *x* · *N*_*e*_ · *r*, where x is the ploidy of the individuals serving as unit count for *N_e_*, and *r* is the recombination rate per base pair (bp) per generation (Dutheil 2024). Both µ and ρ are critical for understanding population trends in species (Santiago et al. 2020)

Mutation rates have conventionally been derived from phylogenies (µ_PHY_), which assume that the rate of substitution accumulation equals the *de novo* mutation rate (Campbell et al. 2021). Estimating µ_PHY_ involves many sources of uncertainty and error, including violations of the molecular clock, incomplete fossil records and dates, and incomplete lineage sorting (Campbell et al. 2021; Suárez-Menéndez et al. 2023). In contrast, parent-offspring trios from pedigrees permit direct estimation of the germline mutation rate (GMR) by identifying *de novo* mutations (DNMs) that arise from replication errors during gametogenesis (Bergeron et al. 2023). The GMR has been quantified across taxa with rates varying from a µ of 10^-11^ in unicellular eukaryotes to a µ of 10^-7^ in multicellular organisms (Bergeron et al. 2023; Suárez-Menéndez et al. 2023). Studies in humans using 3,000 trios estimated µ at ∼1.25 x 10^-8^, while the GMR across a variety of mammals inferred from trios ranged from 3.86 x 10^-9^ to 1.43 x 10^-8^ (Bergeron et al. 2021; Bergeron et al. 2023). Trio-based mutation-rate estimates are sensitive to sequencing depth, with low coverage reducing sensitivity and inflating false-negatives (Wang et al. 2026).

Meiosis and recombination, while highly regulated, are variable across the genome, taxa, sexes, and individuals (Stapley et al. 2017). The distribution of recombination events across the genome is non-random, and the regional variation in recombination rate is associated with genomic features such as GC content and nucleotide diversity (Stapley et al. 2017; Torres et al. 2023). At least one crossover event is required for accurate segregation of homologous chromosomes, and in mammals, there are generally multiple crossovers per chromosome (Hinch et al. 2019; Wooldridge and Dumont 2022). Recombination is also sexually dimorphic, with females having higher average crossover counts than males (Wooldridge and Dumont 2022). Mammalian recombination rates are elevated near telomeres and are suppressed in heterochromatic centromeric regions, with recombination events occurring in hotspots 1 – 5kb long; consequently, most of the genome is recombinationally inert (Wooldridge and Dumont 2022). In most mammals, recombination hotspot locations are defined by the zinc finger protein *PRDM9* (Wooldridge and Dumont 2022).

Polar bears (*Ursus maritimus*) are adapted to the extreme environment and harsh cold of the Arctic (Figure S1A; IUCN (2015)) and rely on the sea ice for feeding, breeding, and in some cases denning (Regehr et al. 2016). Climate-mediated reductions in the availability of sea ice habitat represent the single greatest threat to the long-term conservation of the species (Regehr et al. 2016). This rate of reduction is accelerated as the Arctic is warming at four times the global rate, necessitating the species’ *Vulnerable* (VU) assessment from The IUCN Red List of Threatened Species (IUCN 2015; Regehr et al. 2016; Rantanen et al. 2022).

Historical demographic models for polar bear and closely related species (brown bear; *Ursus arctos*) have been inferred using population genetic methods (Liu et al. 2014). These studies use µ and *r* values that are highly uncertain (µ = 1.83 x 10^-8^; Liu et al. 2014), or borrowed from humans and other mammals (µ = 1 – 1.15 x 10^-8^; Cahill et al. 2013; Kumar et al. 2017; R.J. Wang et al. 2022). Likewise, ursid studies using *r* have used what appear to be arbitrary values (Cahill et al. 2013; de Jong et al. 2023), or values based on domestic dogs (Puckett et al. 2023). Liu et al. (2014) noted that their demographic parameter estimates differed between methods due to reliance on one or both of µ and *r,* and their uncertainty in those rates. Because estimates of *Nₑ* depend on μ and, for linkage-based methods, on *r*, errors in these parameters propagate to any downstream inference or projection that relies on *Nₑ*, an essential parameter in conservation genetics (Mergeay et al. 2026).

In this study, we aimed to produce the first estimates of µ and *r* for the polar bear by estimating a mutation rate from high-coverage whole-genome resequencing data of parent-offspring trios (Figure S1B). Recombination rates and hotspots were estimated from whole-genome resequencing of a polar bear subpopulation. Further, we tested predictions about the variation in *r* between genomic features. Finally, we cross-validated our population-scaled recombination map with pedigree-derived crossover counts from trio phasing. Our findings fill existing gaps in the evolutionary landscape of polar bears.

## 2. Methods

### 2.1 Sample Collection

All individuals were from the Western Hudson Bay (WHB) polar bear subpopulation, sampled through long-term monitoring by Environment and Climate Change Canada (1966-2011; Malenfant et al. 2016; Biddlecombe et al. 2025). Tissue samples were extracted from pinna (external ear) tissue or from a biopsy of superficial rump fat, and blood samples were drawn from the femoral or jugular vein (Newediuk et al. 2025). Genomic DNA was extracted using Qiagen DNeasy Blood & Tissue Kits following the manufacturer’s protocol, as described in (Newediuk et al. 2025).

We identified parent-offspring trios from the WHB genetic pedigree of 4449 individuals (Malenfant et al. 2018). The pedigree has since been updated with additional samples using the same methods and now includes 4,651 individuals with 3,173 maternity and 2,685 paternity assignments. From this pedigree, we selected eight trios (16 individuals) in which both parents and the offspring had archived DNA for high-coverage sequencing. These trios formed three connected pedigree components (Figure S1B). DNA samples were sent to The Centre for Applied Genomics (TCAG) for sequencing at SickKids Hospital (Toronto, Ontario, Canada), where they were sequenced on multiple Illumina NovaSeq X lanes to an average depth of 46x. We confirmed parentage and validated trios based on relatedness estimates from the genome-wide data.

The population sampled for the recombination analysis consisted of 44 individuals from the WHB without any first-order relatives (relatedness phi (ϕ) > 0.177). Samples were selected from long-term monitoring programs, biopsy darting, and hunter harvest submissions, following relevant permit protocols. We extracted DNA from muscle and skin tissue with Qiagen DNeasy Blood & Tissue Kits following the manufacturer’s protocol (QIAGEN®, Germantown, MD, USA). Libraries were prepared and sequenced on Illumina NovaSeq X 25B flow cells with 150 bp paired-end reads and 20X coverage.

All research and handling protocols were approved by Environment and Climate Change Canada’s Western and Northern Animal Care Committee and conducted under research permits issued by Parks Canada and the Province of Manitoba.

### 2.2 Genome Annotation

We extracted exons and genes from the polar bear reference genome GTF file (ASM1731132v1), allowing the inference of introns and intergenic regions using BedTools version 2.31.0 (Quinlan and Hall 2010). We generated a de novo repeat library for the genome using RepeatModeler version 2.0.7 (Flynn et al. 2020), which was used to identify transposable elements (TEs) in the genome assembly with RepeatMasker version 4.2.1 (Smit et al. 2013). We classified RepeatMasker annotations into four TE classes: DNA transposons, LINEs, SINEs, and LTR retrotransposons. For each class, we merged overlapping annotations into non-redundant intervals using BEDTools to generate genome-wide TE class BED files for downstream analyses (Quinlan and Hall 2010). In the GTF file, 5′ UTRs were not annotated; thus, we proxied them using the 100-bp upstream of each gene’s first exon (Torres et al. 2023). We used BEDTools maskfasta to mask annotated exons and TEs. We obtained the GC-base composition for regions with the BEDTools -nuc option and corrected for the masked positions (Quinlan and Hall 2010). We calculated both gene and TE densities as the proportion of a region covered by the annotated sequences of each category. To assess the covariation of each feature with GC, we computed pairwise Spearman’s rank correlations between GC content and each of the TEs and gene density across 1-Mb windows.

### 2.3 Mutation Rate

We used a modified version of the Bergeron et al., 2021 pipeline to detect DNMs. SOAPnuke was used to filter out adaptors, low-quality reads and N-reads (Y. Chen et al. 2018). We mapped the trimmed reads to the *Ursus maritimus* reference genome using BWA-MEM version 0.7.18 (Li 2013). We then removed duplicate alignments with Picard MarkDuplicates 3.0.0. (GATK Team 2024). We completed variant calling using GATK (Genome Analysis Toolkit) version 4.4.0.0 (GATK Team 2024). We called variants using HaplotypeCaller in BP-Resolution mode (GATK Team 2024). We combined all gVCF files per individual into a single file per trio and per scaffold using CombineGVCFs (GATK Team 2024). We then applied joint genotyping with GenotypeGVCF (GATK Team 2024). We then conducted site filtering to detect *de novo* mutations with the following parameters: QD < 2.0, FS > 20.0, MQ < 40.0, MQRankSum < –2.0, MQRankSum > 4.0, ReadPosRankSum < –3.0, ReadPosRankSum > 3.0, and SOR > 3.0. We applied the following filters:

i. We selected Mendelian violations with GATK SelectVariants such that only sites where both parents were homozygous reference (HomRef), and the offspring was a heterozygote (Het);
ii. We used an allelic balance filter for *de novo* mutations to ensure the number of alternative alleles in the offspring accounted for between 30% and 70% of total reads;
iii. We filtered depth per trio to range between 0.5 x m_depth_ and 2 x m_depth_, with m_depth_ being the mean depth of the trio to avoid erroneous calls due to sequencing error; and
iv. We set a genotype quality (GQ) filter at a GQ of 60.

We then split VCF files per trio using GATK SelectVariants (GATK Team 2024). We then used BCFtools version 1.19 to calculate the callability of the trio’s genomes using bcftools view to apply site filters (Danecek et al. 2021). We calculated the false negative rate of each trio with a custom R script that used the callable sites identified by BCFtools after they were converted to a table using GATK VariantsToTable (Danecek et al. 2021; GATK Team 2024).

To prevent the inclusion of false-positive DNMs, we called variants in the region of these putative DNMs using a second variant caller (BCFtools version 1.19; (Danecek et al. 2021)) (Bergeron et al. 2023). We removed false-positive calls, which are sites where at least one parent has the same variant as the offspring, or where the offspring has no variant (Bergeron et al. 2023). This formed a consensus set of DNMs considered true positive calls. Finally, we calculated the number of DNMs using a custom R script.

We estimated the mutation rate per site per generation per trio with the following equation:

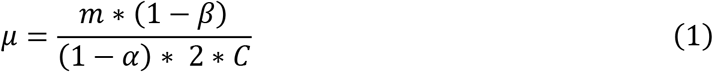

where µ is the mutation rate, *m* is the number of de novo mutations, ß is the false positive rate, α is the normalized false negative rate, and finally, C is the size of the callable genome. Candidate DNMs that we rejected are reported per trio as the false-positive rate in Table 2 and were excluded from the final DNM counts. We applied no further false-positive correction to the final counts (i.e., residual FPR = 0), as the stringent filtering and variant caller concordance make it unlikely that false positives remained in the final dataset <u>(Suárez-Menéndez et al., 2023)</u>. We defined the false negative rate (FNR; α) of a trio in two parts:

i. The proportion of true heterozygote sites outside the allelic balance threshold where one parent is HomRef, the other parent HomAlt, and their offspring Het (FNR_AB_) <u>(Bergeron et al., 2023)</u>.
ii. The proportion of these heterozygous sites, restricted to those passing the GQ and depth filters that were removed by our site filters (FNR_site_) (Bergeron et al. 2023).

We combined the pooled rates as α = 1 − (1 − FNR_AB_) (1 − FNR_site_), which we applied to all trios in Equation 1. Callable genome size *C* is the number of sites where both parents are HomRef (filter i), the three individuals pass both the depth filter (filter iii), and all three individuals pass the quality filter (filter iv). The pipeline is in Figure S2.

### 2.4 Recombination Landscape and Nucleotide Diversity

We processed raw reads from the population sequencing effort (44 WHB bears), and we generated BAM files through the snpArcher v1 workflow (Mirchandani et al. 2024). We preprocessed reads by trimming adapter sequences and reads with greater than 40% of bases with a Phred score below Q15 with fastp v0.20.1 (S. Chen et al. 2018). We aligned the raw reads to the *Ursus maritimus* assembly (Laidre et al. 2022). We aligned sequences to the reference genome using BWA-MEM v0.7.17 (Li 2013) to produce per-sample BAM files, and removed PCR and technical duplicates with sambamba v0.8.0 (Tarasov et al. 2015).

Once BAM files were generated, we departed from the snpArcher workflow. We downsampled the BAM files to 10X coverage to enable even calling across the broader sample set (Rivkin LR personal communication) by subsampling individual BAM files to reduce read counts with samtools v1.13 (Danecek et al. 2021). We created custom UNIX scripts to jointly call SNPs across all samples using bcftools v1.17 (Danecek et al. 2021). Given the size of our dataset, we minimized processing time by calling SNPs separately across scaffolds in 10 Mb intervals. We used bcftools to remove reads with low quality scores, strand bias, read mismatches, and multi-allelic calls, and retained reads with a minimum depth > 10x and genotype quality > 30 in at least one sample per interval (QUAL < 30, DP < 10, SP > 3, SOR > 3.0, MQ < 30.0, GQ < 30). Finally, we removed indels and multi-allelic calls and retained variants (SNPs). Following filtering, we concatenated intervals into a single VCF file that included all samples. The resulting VCF file had 7,773,328 variants, with a genotyping rate of 0.998.

To estimate *r* in the population, we computed the recombination rate using pyrho version 0.2.1 (Kamm et al. 2016; Spence and Song 2019). We first stripped the population VCF file of all INFO metadata, retaining only genotype format fields. We then generated a demographic-aware recombination lookup table using pyrho make_table (Kamm et al. 2016; Spence and Song 2019), incorporating an MSMC-derived polar bear population history (Clendenin et al. 2025), and converted it to absolute population sizes and generations using our pedigree-based mutation rate. We determined optimal model parameters using pyrho hyperparam by simulating a diploid ploidy structure across a grid search of block penalties (10, 20, 50, 100) and window sizes (20, 25, 50, 100) (Kamm et al. 2016; Spence and Song 2019). We then inferred fine-scale recombination maps for each scaffold using pyrho optimize with a selected window size of 100 and a block penalty of 10 (Kamm et al. 2016; Spence and Song 2019). We converted raw outputs to genetic distances (cM/Mb). Following standard practices of removing localized rate spikes indicative of local artifacts, we identified windows with implausibly high inferred rates (*r* (cM/Mb) > 100) (Auton et al. 2012; Pfeifer 2020; Soni et al. 2026). In total, we identified 4,830 such regions which were masked across all scaffolds, along with the 50 SNPs upstream and downstream, and excluded from downstream summaries. Additionally, recombination rates at scaffold boundaries are poorly constrained; thus, we masked the first and last pyrho block of any scaffold whose inferred rate exceeded the 99th percentile of 1-Mb window rates in the initial map. Finally, we summarized rates into fixed 50kb and 1Mb windows using a custom Python script. We used 1-Mb window aggregation only for describing landscape variation and for the feature-association regression.

We estimated the average number of nucleotide differences between sequences drawn from the same population (Nei and Li 1979), nucleotide diversity (π), with pixy version 2.2.3 (Korunes and Samuk 2021) from an all-sites population VCF file in 10kb non-overlapping windows. We aggregated windows to 50kb to match the resolution of the pyrho recombination map. Filtering consisted of two steps: i) excluded windows where π > 5 x 10^-3^, indicative of copy-number variation or collapsed paralogs (Hartasánchez et al. 2018); and ii) excluded windows with low-density windows with < 25kb callable sites (50% of total window size) to ensure sufficient data and no mapping artifacts.

### 2.5 Recombination Hotspots

From the population VCF, we inferred genome-wide fine-scale recombination rates to detect hotspots in polar bears using LDhat version 2.2 (McVean et al. 2002; McVean et al. 2004; Auton and McVean 2007). We generated a lookup table containing the coalescent likelihoods for every two-locus haplotype configuration possible in our sample of 44 diploid individuals (i.e., 88 haploids: -n 88), using LDhat complete with the suggested maximum population-scaled recombination rate ρ of 100 (-rhomax 100) and a grid size of 101 (-n_pts 101). We then derived estimates by running LDhat’s interval function with a block penalty of 5 (-bpen 5) for 60 million iterations (-its 60000000) with a sampling scheme of 40,000 iterations (-samp 40000) across each scaffold. We used LDhat’s stat function to discard the burn-in of the MCMC using (-burn 500). Once again, following standard practices of removing localized rate spikes, we identified windows with implausibly high inferred rates (ρ > 100) (Auton et al. 2012; Pfeifer 2020; Soni et al. 2026).

We inferred recombination hotspots using LDHot version 0.4, a statistical method that directly processes the recombination rate estimates obtained from LDhat (Auton et al. 2014). First, we ran ldhot to identify putative recombination hotspots using the default parameters (3 kb windows, 1kb step size, background window 100 kb, and 1,000 simulations) (Auton et al. 2014). Next, we merged significant windows using ldhot_summary with significance thresholds of 0.001 and 0.01 for calling and merging hotspots, respectively.

To filter the putative hotspots identified by LDHot and obtain a high-quality dataset, we applied a three-step filtering approach assembled by Soni et al. (2026), which combines recommendations from other studies (Stevison et al. 2016; Brazier and Glémin 2024). We excluded hotspot candidates wider than 10 kb, as recombination hotspots are generally narrow. Artificial breaks in LD resulting from mis-assembly or mis-alignment can lead to localized peaks in recombination rate. We calculated the intensity of each hotspot candidate by dividing the peak recombination rate by the rate of the surrounding 100 kb region and removed candidates with an intensity higher than 200. Additionally, we applied a low-intensity filter to exclude candidates with an intensity lower than four due to the difficulty of distinguishing genuine low-intensity hotspots from background variation. Finally, to account for recombination rate variation across chromosomes and further narrow down the location of recombination hotspots, we divided each remaining candidate region into 2kb windows with a step size of 1 kb, keeping only the hotspot windows exhibiting rates of at least five times the chromosome-wide average rate.

### 2.6 Recombination Breakpoints

We identified crossover breakpoints directly from our parent-offspring trios using pedigree-aware phasing with whatshap version 2.8 (Martin et al. 2016). We phased the VCF files using the whatshap phase function; we provided the tool with the joint-genotyped VCF file per scaffold, along with the pedigree using the -ped mode (Garg et al. 2016; Martin et al. 2016). We called crossovers from the phased VCFs with a custom Python script that scans, per meiosis, for transitions between parental haplotypes in a sliding window of informative sites. The script requires a run of at least three consecutive informative sites supporting a new parental-haplotype assignment before accepting a switch (isolated discordances are treated as genotyping noise). We cancelled pairs of switches that reverted to their original state within 1 Mb of each other as local phasing artifacts rather than counted as double recombinants. To reduce spurious breakpoints attributed to mis-mapping in repetitive regions, we removed crossovers overlapping any repeat/TE annotation from the BED files (see "Genome Annotation” above). The remaining set formed our pedigree-derived crossovers, which we used to summarize per-meiosis crossover counts, estimate the pedigree-based recombination rate, and to validate the pyrho and LDhot rate estimates. We calculated the sex-averaged autosomal recombination rate as the mean number of crossovers per meiosis (equivalent to map length in Morgans) x 100 (equaling cM), divided by the total length (Mb) of the autosomal genome analyzed (reference genome ∼2175 Mb).

### 2.7 Enrichment and Genomic Features

To test whether the pedigree-derived crossovers and our trio-based DNMs coincide with recombination hotspots identified independently from population-level linkage disequilibrium, we performed a permutation-based enrichment test. For each crossover breakpoint set, we counted the number of breakpoints overlapping an LDhot hotspot interval. We then generated a null distribution by shuffling the positions of the observed crossover breakpoints 10,000 times using bedtools shuffle (Quinlan and Hall 2010). This was restricted to the same non-repeat-masked genomic space used for crossover calling. We used the same process for DNMs. We calculated fold-enrichment as the observed overlap count divided by the mean overlap count across permutations. We calculated an empirical p-value as the proportion of permutations with an overlap count at least as extreme as the observed value.

To assess the variation in recombination rate among genomic features, we used a linear regression model using the lm function in base R (R Core Team 2021) with recombination rate (1Mb scale) as the response variable. The explanatory variables were GC content, gene density, and the relative abundance of four TE classes (DNA transposons, LINE, SINE, and LTR retrotransposons). We also analyzed the associations between the recombination rate and genomic features using Spearman’s rank correlation tests with the corr option in base R (R Core Team 2021).

To evaluate variation in recombination rates across genomic features, we estimated recombination rates within each TE class and across gene regions (exons, introns, upstream (5’UTR), and intergenic regions). To avoid biases caused by differential selective pressures, we considered the intervals between SNPs (identified by pyrho) within a feature. For example, for a given exon, if an interval between markers overlapped completely with the predicted exon, we retained it, and if the overlap was only partial (including part of the exon sequence but also part of a neighbouring intron or UTR), we discarded it (Torres et al. 2023). These overlaps were identified with BEDTools intersect (Quinlan and Hall 2010).

## 3. Results

### 3.1 De novo mutations

The mean coverage was 46x (minimum 16x, maximum 54x) (SE = 2.07), with a mean breadth at 20x coverage of 89.84% (SE = 0.05%). The trios consisted of 3 unrelated families, with several full and half-siblings (Figure 1; Relatedness phi (ϕ) in Figure S3). An average of ∼43 DNMs per trio (SE = 3.75) passed the initial filters. After calling putative *de novo* mutations with a second variant caller, 329 sites were considered true-positive calls, averaging ∼41 per trio (Table 1). The number of callable sites ranged from 484,663,860 to 2,062,756,345 base pairs, covering an average of 80% of autosomal sites of the polar bear genome. The false-negative rate was 2.79%. Collectively across trios, this yielded a mean PB mutation rate of 1.22 x 10^-8^ *de novo* mutations per site per generation (95% CI of the mean = 0.937 x 10^-8^ to 1.51 x 10^-8^).

**Figure 1.**
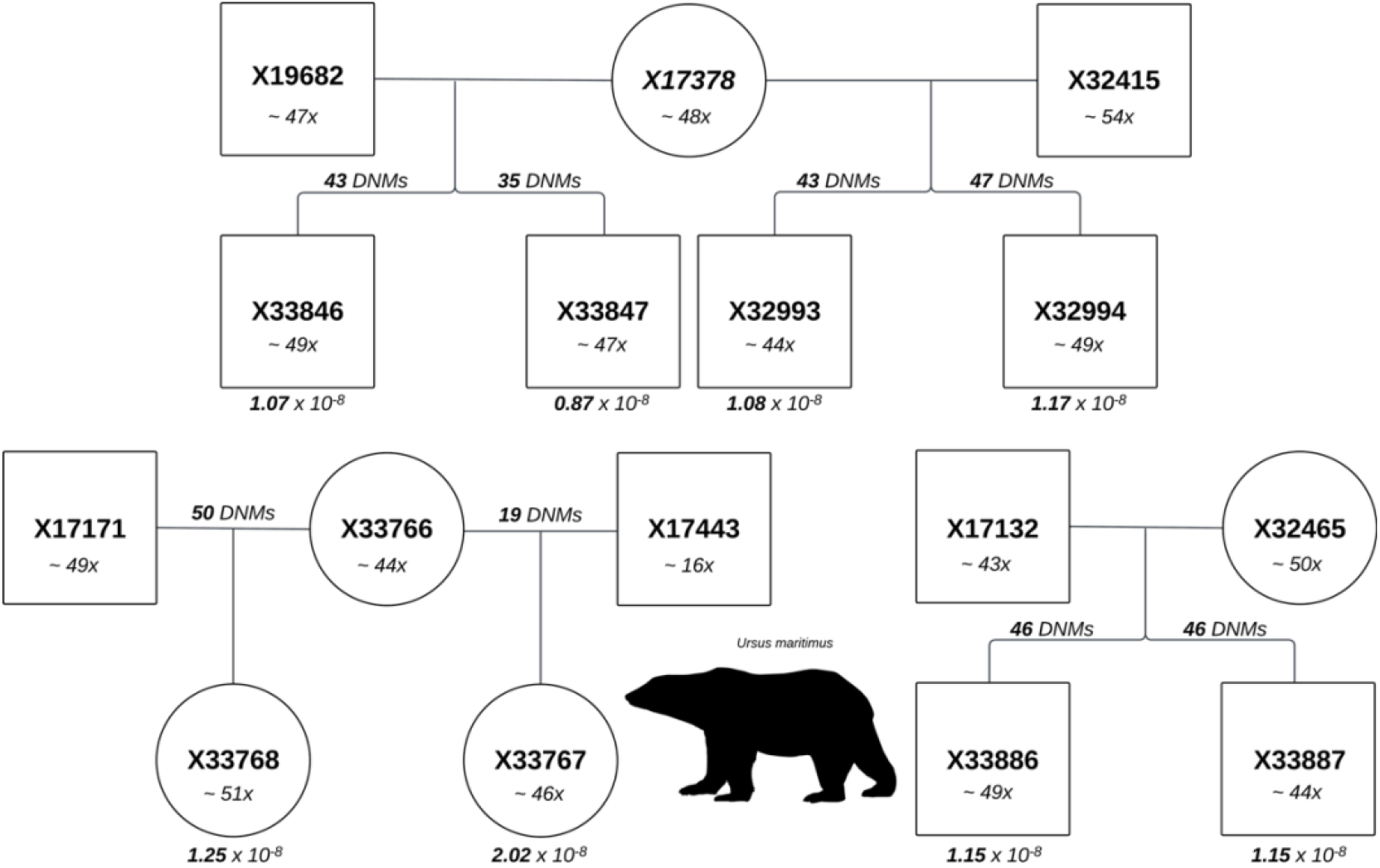
Pedigree of parent-offspring trios used to estimate the germline mutation rate. Each individual: Sample ID (top), coverage (below), DNMs (branches), and mutation rate (bottom).

**Table 1.** Parent-Offspring Trio Mutation Rate. µPED calculated with Equation 1 using the pooled FNR (α = 0.0279)

| **Trio** | **Callable Genome (bp)** | **FNR (allele balance)** | **Initial DNMs** | **Final DNMs** | **FPR** | **µPED** |
|---|---|---|---|---|---|---|
| X32993 | 2,053,971,849 | 0.017 | 45 | 43 | 0.044 | 1.08 x 10<sup>-8</sup> |
| X32994 | 2,062,580,728 | 0.013 | 47 | 47 | 0.000 | 1.17 x 10<sup>-8</sup> |
| X33767 | 484,663,860 | 0.012 | 19 | 19 | 0.000 | 2.02 x 10<sup>-8</sup> |
| X33768 | 2,052,481,195 | 0.011 | 52 | 50 | 0.038 | 1.25 x 10<sup>-8</sup> |
| X33846 | 2,062,756,345 | 0.012 | 44 | 43 | 0.023 | 1.07 x 10<sup>-8</sup> |
| X33847 | 2,058,421,393 | 0.014 | 36 | 35 | 0.028 | 0.87 x 10<sup>-8</sup> |
| X33886 | 2,053,691,929 | 0.012 | 49 | 46 | 0.061 | 1.15 x 10<sup>-8</sup> |
| X33887 | 2,051,401,277 | 0.016 | 48 | 46 | 0.042 | 1.15 x 10<sup>-8</sup> |
| **Average** | **1,859,996,072** | **0.013** | **42.5** | **41.1** | **0.030** | **1.22 x 10<sup>-8</sup>** |

### 3.2 Recombination Landscape

The genome-wide recombination rate estimated by pyrho across the 36 autosomal scaffolds of ASM1731132v1, computed as a length-weighted mean over per-interval rates, was 3.03 cM/Mb with a length-weighted median of 1.28 cM/Mb. Aggregation of pyrho intervals into 1-Mb windows (Figure 2) shifted the mean and median up to 3.06 cM/Mb and 2.78 cM/Mb, respectively, reflecting the smoothing of within-window rate variance across a heavily right-skewed distribution. Recombination rates varied ∼8-fold among full-length 1-Mb windows (n = 2,155; 1^st^ – 99^th^ percentile: 0.98 – 7.62 cM/Mb; range 0.43 – 11.79 cM/Mb), and the top 5% of windows contributed 11.2% of total map length, more than twice the expectation under uniform rates. Per-Mb recombination rates were strongly negatively correlated with scaffold length (Spearman’s ρ_s_ = −0.76, p = 5.49 x 10⁻⁷). All scaffolds in the assembly used for our analysis were autosomal; chromosome quotient values ranged from 0.991 to 1.004 (se=4.89 x 10^-4^) (Table S1). The genome-wide nucleotide diversity (π) was 4.43 x 10^-4^ (sd = 7.91 x 10^-4^), ranging from 2.27 x 10^-6^ to 4.75 x 10^-2^. After filtering out potential CNV/paralog windows (π > 5e-3), nucleotide diversity was positively rank-correlated (non-parametric) with recombination rate (Spearman’s ρ_s_ = 0.11, P < 2.2 x 10^-16^) (Figure S4A). Binning windows into 20 bins by *r* quantile revealed a large reduction in mean π in the lowest-r bin (3.06 x 10⁻⁴ in the bottom 5% of r vs. plateau mean of 4.21 x 10⁻⁴ across bins 2 – 19), a ∼27% reduction, with a similar median π reduction of ∼28% (2.33 x 10⁻⁴ vs. plateau median 3.26 x 10⁻⁴; Figure S4B).

**Figure 2.**
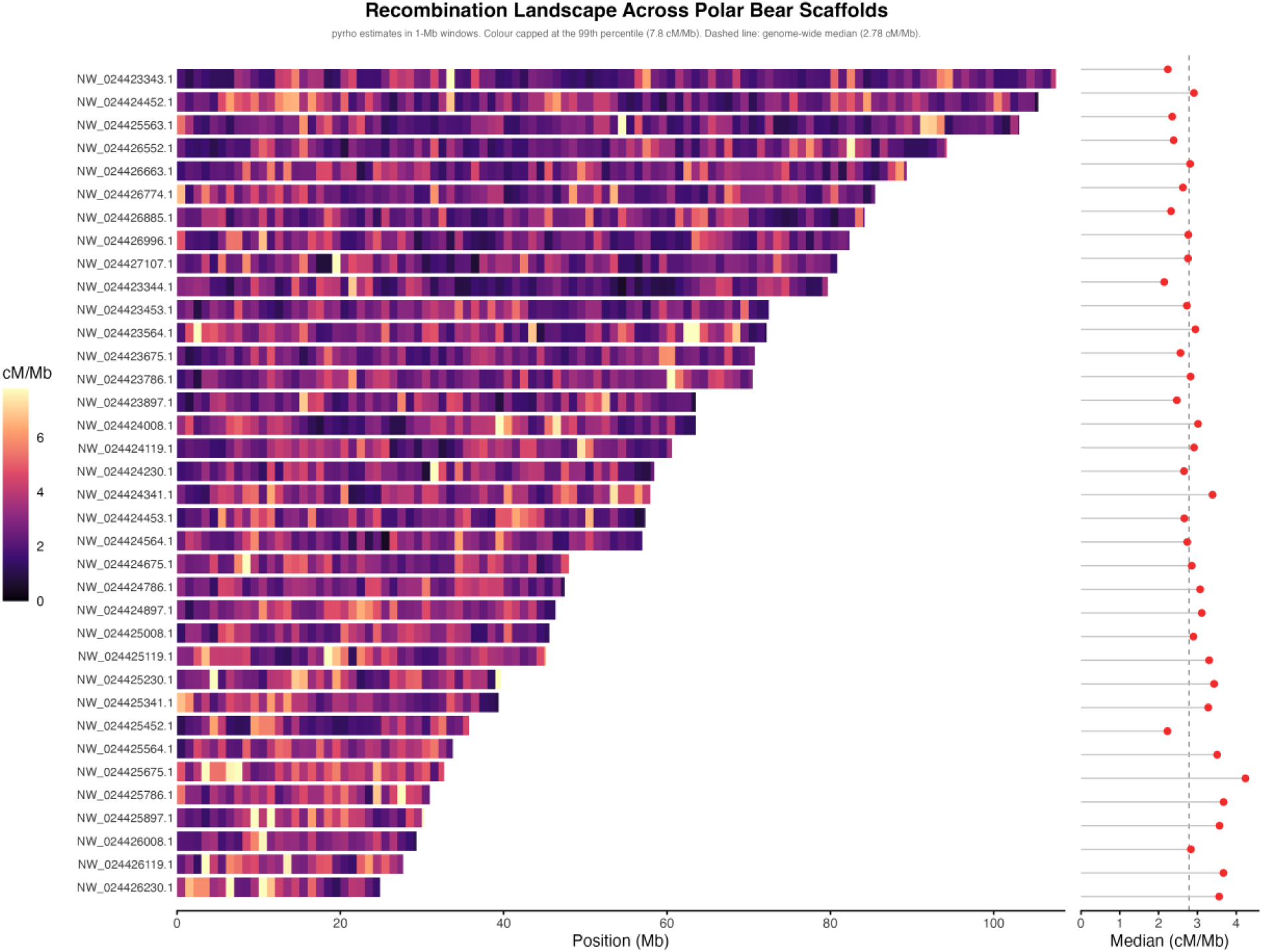
Estimates of the recombination rate (cM/Mb) in 1-Mb windows, chromosomes ordered by chromosome length.

### 3.3 Recombination Hotspot Identification and Enrichment

After filtering, we identified 38,417 candidate hotspot windows (2kb), forming 8,515 distinct hotspot regions that cover 2.10% of the genome. The mean hotspot width was 5.36 kb, with a median of 4.00 kb. Hotspot regions exhibited a median peak-rate-to-background intensity of 18.7x (IQR 9.8 – 35.7) relative to the surrounding 100 kb; typically, a hotspot exceeded its flanking 100-kb baseline by more than an order of magnitude. Although hotspots covered only 2.10% of the genome, they accounted for 43.8% of the total inferred genetic map length across all intervals (the sum of rate multiplied by width), concentrating recombination into hotspot intervals by 20.9-fold. Pedigree-derived crossovers overlapped LDhot-identified hotspots (8,515 regions spanning 2.1% of the genome) more often than expected by chance; of the 522 crossovers identified across all 16 meioses, 47 (9.0%) fell within a hotspot interval, against a null mean of 9.3 (1.8%) across 10,000 permutations, reflecting a 5.06-fold enrichment (empirical p < 0.001). Restricting the analysis to the 14 meioses excluding the two flagged transmissions gave a consistent result: 35 of 396 crossovers (8.8%) overlapped a hotspot, against a null mean of 7.2 (1.8%; 4.84-fold enrichment, p < 0.001). Pedigree DNMs overlapped LDhot-identified hotspots at a similar rate to what would be expected by chance; the 329 DNMs identified across all 8 parent-offspring trios fell within a hotspot interval 6 times (1.8%), against a null mean of 5.12 (1.6%) across 10,000 permutations, a 1.17-fold enrichment (empirical p = 0.41).

### 3.4 Recombination Breakpoints

Trio-based phasing across all 36 scaffolds and 16 meioses yielded ∼250 candidate crossovers per meiosis before repeat-based filtering. After removing crossovers that overlapped annotated repeat/TE intervals from the class-wise BED files, 522 high-confidence pedigree crossovers survived across the 16 meioses (mean 32.6 crossovers per meiosis), which is within the range reported for other mammals. This yielded per-meiosis counts that were generally consistent, with two notable exceptions: X33767’s paternal meiosis (father X17443) was elevated (57 crossovers), and X33768’s paternal meiosis (father X17171) was the highest in the dataset (69 crossovers). Because each crossover contributes one Morgan of map length per gamete, this implies a sex-averaged autosomal map length of ∼3,260 cM and a genome-wide recombination rate of ∼1.50 cM/Mb (1.30 cM/Mb excluding the two elevated paternal meioses). Crossover density was negatively correlated with scaffold length (r = -0.52).

### 3.5 Associations between recombination rate and genomic features

The model revealed a significant association between the recombination rate and the explanatory variables (F = 12.19, df = 6 and 2184, P = 1.58 x 10^-13^) yet explained only a marginal amount of the total variation (R^2^ = 3.24 x 10^−2^, Adj R^2^ = 2.98 x 10^−2^). Only GC and SINEs were significant (α = 0.05) among the explanatory variables (Table 2).

**Table 2.**
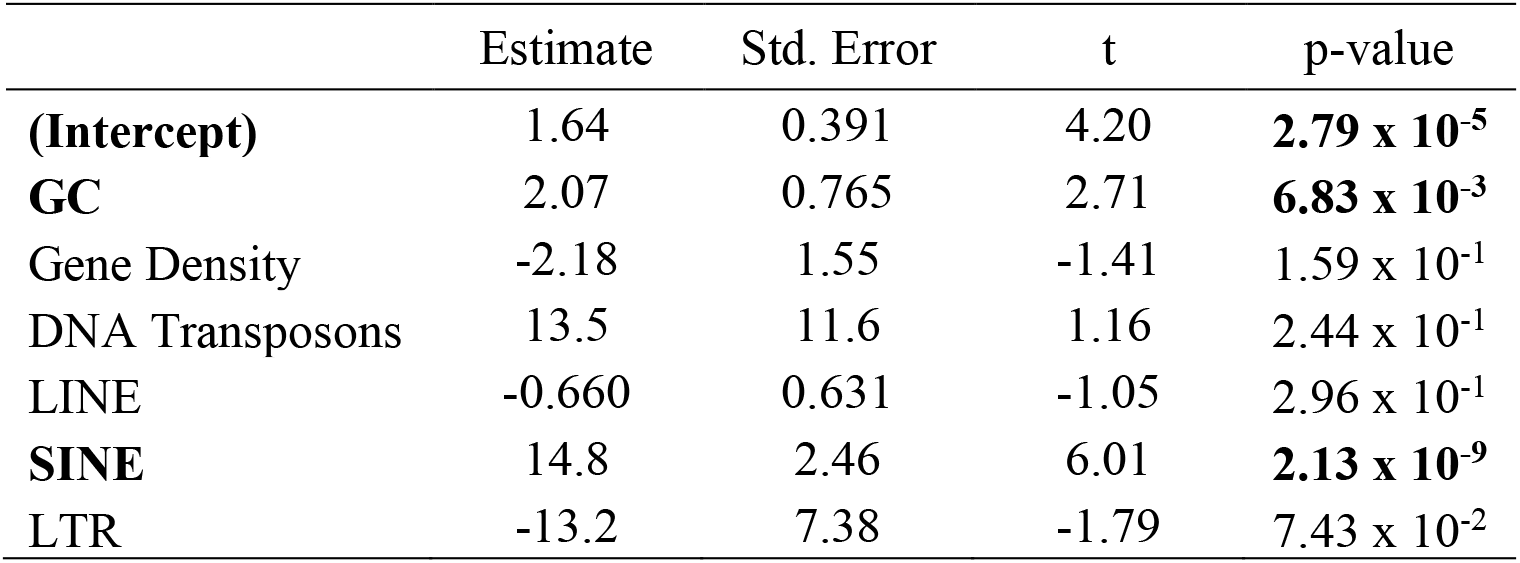
Summary of the results from a linear regression model. Explanatory variables significantly associated with recombination rate variation at a 1-Mb scale are highlighted in bold.

The strength and direction of the recombination rate were differently associated between TEs. SINEs (Spearman’s ρ_s_ = 0.16, P = 2.47 x 10^-13^, Fig. S5C) were positively associated, whereas LTRs (ρ_s_ = −0.08, P = 8.59 x 10^-5^, Fig. S5B) and LINEs (ρ_s_ = −0.12, P = 1.43 x 10^-8^, Fig. S5D) were negatively associated with the recombination rate and DNA transposons showed no significant association (ρ_s_ = 0.03, P = 0.11, Fig. S5A). The strength and direction of feature-*r* associations largely mirrored each feature’s covariation with GC content (Fig. S6): gene density (ρ_s_ = 0.64) and SINEs (ρ_s_ = 0.27) were positively correlated with GC, while LINEs (ρ_s_ = −0.44), LTRs (ρ_s_ = −0.33), and DNA transposons (ρ_s_ = −0.19) were negatively correlated (all p-values < 2.2 x 10⁻¹⁶).

We computed length-weighted quantile summaries of the pyrho rate map restricted to intervals falling wholly within each feature category (Figure 3). Relative to the length-weighted genome-wide mean recombination rate of 3.03 cM/Mb (median 1.28 cM/Mb), the strongest local elevation was observed in the 100-bp region immediately upstream of gene starts, the 5′ UTR proxy, at a median of 1.82 cM/Mb (IQR 0.95 – 3.72). Exons (median 1.42, IQR 0.72 – 2.68), SINEs (median 1.45, IQR 0.75 – 2.68), and LINEs (median 1.41, IQR 0.74 – 2.74) were also elevated, introns were slightly above the genome-wide median (median 1.31, IQR 0.68 – 2.46), while intergenic (median 1.26, IQR 0.65 – 2.37) and DNA-transposon (median 1.25, IQR 0.62 – 2.40) intervals were approximately at the genome-wide median. LTRs showed the greatest reduction in local recombination rate (median 1.18 cM/Mb, IQR 0.61 – 2.16), matching the negative direction of the LTR-*r* correlation reported above. LINE-overlapping intervals had an elevated median rate despite the negative association between LINE density and recombination rate at the 1-Mb scale, suggesting that the LINE-*r* relationship is scale-dependent. DNM-overlapping intervals had a median recombination rate of 1.44 cM/Mb (IQR 0.71 – 2.63), ∼13% above the genome-wide median, suggesting a small tendency for de novo mutations to fall in higher-recombination regions.

**Figure 3.**
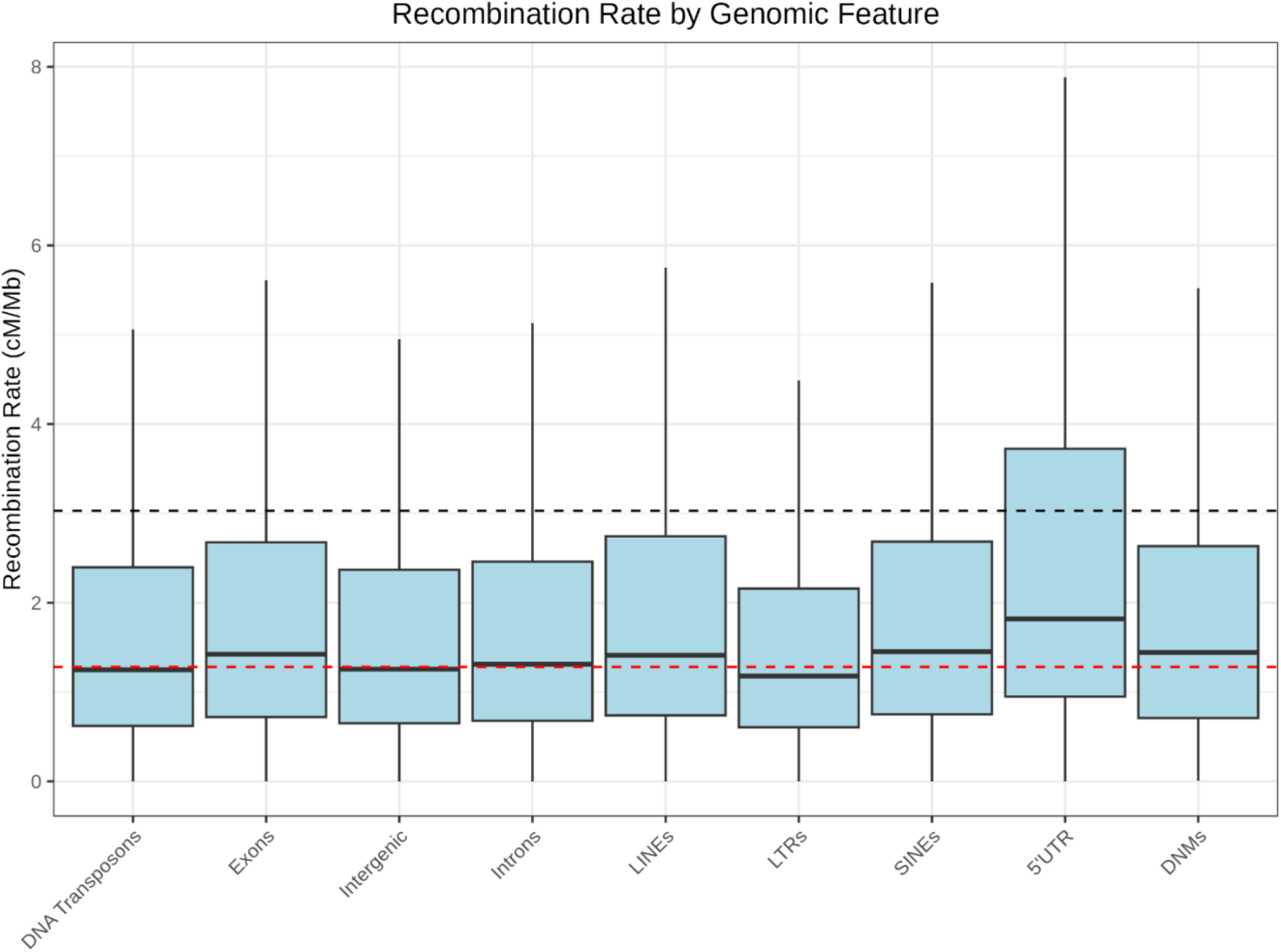
Recombination rate estimates in different genomic features. The mean genome-wide recombination rate (3.03 cM/Mb; black dashed line) and the median genome-wide recombination rate (1.28 cM/Mb; red dashed line) are indicated.

## 4. Discussion

We generated the first pedigree-based estimates of the mutation rate (μ) and the first recombination rate maps for the polar bear. This work addresses a long-standing gap in which demographic and conservation genomic inference for this species has relied on μ and *r* values borrowed from humans or other mammals rather than estimated directly (Cahill et al. 2013; Liu et al. 2014; Kumar et al. 2017; R.J. Wang et al. 2022; de Jong et al. 2023). Our results provide four species-specific parameter sets: a pedigree-based mutation rate μPED, a demographic-aware recombination rate, a hotspot catalogue, and pedigree-derived crossovers and their associated recombination rate (1.50 cM/Mb). Together, these clarify how the polar bear recombination landscape aligns with and diverges from typical mammalian expectations and can be used to refine future demographic reconstructions and climate-linked population projections for this IUCN Vulnerable, ice-obligate species (IUCN 2015; Regehr et al. 2016; Rantanen et al. 2022).

### 4.1 De Novo Mutations

Our μPED estimate of 1.22 x 10^-8^ mutations per site per generation sits within the mammalian range reported across trio-based studies (3.86 x 10^-9^ – 1.43 x 10^-8^; Bergeron et al. 2021; Bergeron et al. 2023), and below the commonly cited human estimate (∼1.25 x 10^-8^; Bergeron et al. 2023). It marginally exceeds the values previously used for polar bear demographic modelling (1 – 1.15 x 10^-8^; Cahill et al. 2013; Kumar et al. 2017; R.J. Wang et al. 2022) and falls within the same order of magnitude as the pedigree-based grizzly bear estimate (µPED = 0.84 x 10^-8^; R.J. Wang et al. 2022).

Several biological factors warrant consideration in our estimate. Parental age at reproduction is a driver of DNM accumulation in mammals (Rahbari et al. 2016; Bergeron et al. 2023), and paternal age scales with DNM count in large human trio datasets (Jónsson et al. 2017). Parental ages were not incorporated into the per-trio estimates we reported, and if the WHB polar bear cohort we sampled skews older relative to the mammalian baseline cohorts, part of our rate could reflect a polar bear-specific age effect. Sequence coverage heterogeneity between trios warranted a check, as sequencing depth is crucial for reducing false-negative and false-positive rates (Wang et al. 2026). Specifically, trio X33767 (father X17443 at ∼16x coverage, versus ∼43x – 54x across the other trios) has the smallest callable genome and the highest estimated μPED. Removing this trio from the pooled mean lowered the estimate to ∼1.11 x 10^-8^ per site per generation (95% CI = 0.997 x 10^-8^ to 1.22 x 10^-8^), a ∼9% reduction. A formal Grubbs test at α = 0.05 confirmed that X33767 is an outlier (G = 2.34 vs G_Crit_ = 2.03), so reporting the estimate excluding the trio with this value (1.11 x 10^-8^) alongside the eight-trio value (1.22 x 10^-8^) is advised. Notably, the value with the exclusion of 1.11 x 10^-8^ falls within the 1.0 – 1.15 x 10^-8^ range previously assumed for polar bear demographic modelling (Cahill et al. 2013; Kumar et al. 2017; M.-S. Wang et al. 2022), suggesting those borrowed values were not far off.

### 4.2 Recombination Landscape and Nucleotide Diversity

The population recombination map yielded a genome-wide *r* of 3.03 cM/Mb (mean) and 1.28 cM/Mb (median) across the 36 autosomal scaffolds of the polar bear reference genome (Figure 2), with the median rate falling close to rates used for ursids (de Jong et al. 2023; Puckett et al. 2023). Aggregating pyrho intervals into 1-Mb windows shifted the median up to 2.78 cM/Mb, a smoothing artifact of the right-skewed distribution expected for recombination rates (Zedek et al. 2026). Rates vary ∼8-fold across the middle 98% of windows (1^st^ – 99^th^ percentile: 0.98 – 7.62 cM/Mb) and the top 5% of windows contributed ∼11.2% of the total map length, more than twice the uniform expectation, and consistent with the concentration of recombination into a small fraction of the genome that has been reported in humans (Halldorsson et al. 2019), house mouse (Wooldridge and Dumont 2022), and dogs (Auton et al. 2013). Per-Mb rates declined with scaffold length, matching the obligate crossover expectation shared across mammalian karyotypes (Jensen-Seaman et al. 2004).

Nucleotide diversity was low across the genome (π = 4.43 x 10^-4^, SD 7.91 x 10^-4^), relative to mammalian averages (2.48 x 10^-3^; Leffler et al. 2012), which matches the bottlenecked demographic history reported for polar bears (Liu et al. 2014), and is consistent with the low genetic diversity expected in carnivores and large mammals (Jeon et al. 2024). This rate was also lower in WHB bears relative to the other subpopulations (4.94 x 10^-4^; Rivkin LR personal communication). Across 42,143 filtered 50-kb windows, we detected a weak but significant relationship between π and *r*. The positive rank correlation indicated a generally monotonic but non-linear π-r relationship which is largely driven by the low-*r* tail, as the mean π in the bottom 5% of *r*-quantile windows was ∼27% lower than the majority of bins (2-19). This trend is predicted by linked-selection theory (Josephs and Wright 2016), where there is reduced diversity in low-recombination regions where the effects of background selection and hitchhiking are strongest (Cutter and Payseur 2013), and its presence in polar bear despite the population’s low overall baseline π shows that in a species whose diversity is reduced by demographic history, recombination-linked variation in π is still detectable at genome-wide scale.

### 4.3 Recombination Breakpoints

Our pedigree-derived crossover catalogue (∼33/meiosis) sits within the range reported for other mammals and is comparable to counts in humans (∼26 in males, ∼44 in females, mean ∼36; Halldorsson et al. 2019). The ∼5-fold enrichment of pedigree crossovers in LDhot hotspots supports our approach capturing genuine biological recombination rather than an artifact of the LD estimator. Per-meiosis counts were broadly consistent across pedigrees, with two elevated meioses: X33767’s paternal meiosis (father X17443, 57 crossovers) and X33768’s paternal meiosis (father X17171, 69 crossovers). The X17443 elevation could be explained by low coverage (∼17x) inflating apparent breakpoint density; however, the X17171 elevation persists at normal coverage (∼49x), and its cause remains uncertain. Despite the increased counts, excluding both meioses did not change hotspot enrichment.

The crossover-derived mean genome-wide recombination rate of ∼1.50 cM/Mb is, to our knowledge, the first direct, pedigree-based estimate of recombination rate in a wild ursid. The pedigree estimate also provides an independent benchmark for our LD-based map. The genome-wide median pyrho rate (1.28 cM/Mb) was comparable to our pedigree rate, and almost identical when excluding the elevated meiosis (1.30 cM/Mb). This agreement is notable given that LD-based rates reflect recombination averaged over many generations and are converted to per-generation rates through our µ value (1.22 x 10^-8^) and integrated the demographic history inferred from Clendenin et al. (2025). Our pedigree-based crossover average recombination rate is broadly comparable to the sex-averaged human rate (∼1.13 cM/Mb; Dumont and Payseur 2008) and sits at the upper end of the mammalian range (0.184 – 1.554 cM/Mb) reported across 13 species (Dumont and Payseur 2008). It is also notable that the polar bear map length (∼3260 cM) falls between those of the dog (∼2,100 cM; Wong et al. 2010) and the domestic cat (∼4,500 cM; Li et al. 2016), and is in line with humans (∼3400 cM; Halldorsson et al. 2019).

### 4.4 Recombination Hotspot Identification and Enrichment

We identified 38,417 candidate recombination hotspot windows (2 kb) that collapsed into 8,515 distinct hotspot regions. These regions accounted for 43.8% of the total inferred map length, a 20.9-fold concentration of recombination into hotspot intervals, indicating that historical recombination in polar bears is highly localized. Another comparison comes from the direct overlap between our LD-defined hotspots and pedigree crossovers; 9.0% of the 522 pedigree crossovers fell within LDhot hotspots, compared to the 71 – 75% of pedigree crossovers falling within hotspots in humans (Halldorsson et al. 2019). This equates to a ∼5-fold enrichment relative to random chance. The concordance of the signal between these two independent approaches provides cross-validation of our LD-inferred hotspot map. To our knowledge, this also marks the first pedigree-vs-LD comparison in a wild mammal.

We also found that DNMs did not overlap LDhot hotspots more than expected by chance (1.17-fold enrichment, empirical p = 0.41). There is a strong DNM-crossover co-localization reported in humans where the DNM rate is elevated up to ∼50-fold within 1 kb of crossovers (Halldorsson et al. 2019). This absence, combined with the slightly above-average median recombination rate for DNM-overlapping intervals (1.44 cM/Mb, ∼13% above the genome-wide median), suggests DNMs in polar bear fall in slightly warmer regions overall but do not concentrate in the narrow hotspot intervals. While neither of these results showed statistical significance, it is worth noting that both results trend in the expected direction with recombination and mutation co-occurring.

A weakened or *PRDM9*-independent hotspot mechanism in Ursidae is an interesting interpretation of the combined pattern of strong promoter-proximal (5’UTR) recombination rates (Figure 3), which parallel the dog landscape (Auton et al. 2013), weak crossover concentration in hotspots, and the null DNM-hotspot enrichment. However, several pieces of evidence argue against this theory. Canids remain the only Caniformia (suborder of Carnivora) branch with documented loss of *PRDM9* zinc-finger binding specificity, whereas the giant panda, an ursid much more closely related to the polar bear, carries a functional *PRDM9* (Axelsson et al. 2012). Additionally, polar-grizzly hybrids produce fertile offspring and show evidence of introgression (Hailer 2015; Shafer 2023), which would be implausible with incompatible or non-functional *PRDM9* in one of the species (Oliver et al. 2009; Mukaj et al. 2020). We can therefore more likely interpret the promoter-proximal enrichment and weak pedigree-hotspot concordance as reflecting the resolution of our LD-and pedigree-based maps rather than a departure from the traditional mammalian *PRDM9* hotspot pathway. Direct characterization of *PRDM9* sequence and expression in polar bear could help resolve this.

### 4.5 Associations Between Recombination Rate and Genomic Features

Our multiple regression of recombination rate on six genomic-feature covariates (GC content, gene density, and the abundance of four TE classes) was significant overall but explained only ∼3.2% of variance in per-window rates. This significance, paired with the small effect size, indicates that at the 1-Mb scale, these features detect real but modest patterns across the polar bear recombination landscape. The overall pattern of positive associations of recombination rate with DNA transposons (non-significant) and SINE abundance, negative associations with LINE and LTR densities, and only a small fraction of per-window variance explained at the 1-Mb scale, largely mirrors what has been reported in mammals and other species (Jensen-Seaman et al. 2004; Tortereau et al. 2012; Torres et al. 2023). Importantly, each TE class’s correlation with recombination rate largely matched its covariation with GC content (Fig. S6), consistent with the mammalian pattern of SINEs accumulating in GC-rich, open-chromatin regions, while LINEs and LTRs preferentially occupy GC-poor isochores (Lander et al. 2001). The TE signals in our regression are therefore likely at least partly mediated by the underlying GC landscape rather than direct effects of the elements themselves.

The within-feature analysis complements this picture. Consistent with the promoter-proximal recombination pattern discussed above, our 5′ UTR proxy showed the highest median recombination rate, and LTR intervals the lowest, matching the negative direction of the LTR association at 1-Mb scale. LINE-overlapping intervals, however, showed a modestly elevated median *r* (1.41 cM/Mb) despite the negative LINE-*r* Spearman correlation at the 1-Mb scale, indicating that the LINE-*r* relationship is scale-dependent. LINE-rich 1-Mb windows tend to have lower recombination on average, but the pyrho intervals that fall wholly within individual LINE elements themselves do not. This scale-dependent signal has been reported in other mammalian genomes where TE-based structure at Mb scale reflects large-scale GC distribution rather than local-level biology (Jensen-Seaman et al. 2004), consistent with our regression analysis. DNM-overlapping intervals had a median *r* of 1.44 cM/Mb, ∼13% above the genome-wide median, a small but consistent tendency for de novo mutations to fall in higher-recombination regions, though weaker than the strong DNM-crossover co-localization reported in humans (Halldorsson et al. 2019) and consistent with the null hotspot-enrichment test.

### 4.6 Broader Implications

The fine-scale recombination landscape demonstrated a positive π-*r* association consistent with linked selection. This result, combined with the strong promoter-proximal recombination enrichment we observed, presents a future test: targeted characterization of *PRDM9* sequence and expression in polar bear and other ursids. This would allow us to distinguish between a limit in our data and a genuine departure from the traditional mammalian hotspot pathway.

This study provides direct, pedigree- and population-based estimates of mutation (μ) and recombination rate (*r*, ρ) for polar bears, replacing values previously borrowed from distantly related taxa in demographic reconstructions of this species (Cahill et al. 2013; Liu et al. 2014; Kumar et al. 2017; R.J. Wang et al. 2022; de Jong et al. 2023). Coalescent-based demographic inference depends directly on μ to scale time and effective population size (Schiffels and Wang 2020) and on *r*/ρ to define the linkage structure over which selection and drift operate (Stapley et al. 2017; Santiago et al. 2020). Thus, the species-specific estimates presented here should improve the accuracy of historical *Nₑ* reconstructions and, by extension, projections of polar bear population trajectories under ongoing Arctic sea ice loss (Rantanen et al. 2022; Biddlecombe et al. 2025).

## Acknowledgements

We thank Environment and Climate Change Canada for sampling.

## Code availability

The code supporting this work is available in our lab’s GitLab repository: https://gitlab.com/WiDGeT_TrentU/

## Data availability

Raw paired-end Illumina whole-genome sequencing reads for the polar bears have been deposited in the NCBI Sequence Read Archive under BioProject PRJNA1556893. The data are under embargo until publication. Reviewers can view the project and sample metadata through a private reviewer link, which will be provided as soon as NCBI provides it. Once released, the data will be available at https://www.ncbi.nlm.nih.gov/bioproject/PRJNA1556893.

## Author Contribution

AS and DD conceived of the study; DD conducted all the analyses. LN and RR conducted lab work. ER and JN collected samples. All authors reviewed and commented on the manuscript.

